# Osteoblasts Exert a Pro-Tumorigenic Effect on Breast Cancer Spheroids Through CXCL5/CXCR2 Signaling In 2D And 3D Bone Mimetic Cultures

**DOI:** 10.1101/2025.11.26.690859

**Authors:** Sarah Nano, Syeda M. Naqvi, Isabella Weiner, Natalie Volz, Vatsal Kumar, Laurie E. Littlepage, Laoise M. McNamara, Glen L. Niebur

## Abstract

Bone provides a favorable niche for breast cancer colonization and metastatic progression. Breast cancer cells are attracted to the bone microenvironment where they induce bone cells to resorb bone, which enhances tumor cell proliferation in a positive feedback loop often referred to as the vicious cycle. While this phenomenon is established, the molecular interactions between cancer cells and bone cells are not well defined. CXCL5/CXCR2 signaling has recently been shown to promote breast cancer colonization to the bone. Here, we investigate the effects of osteoblasts and osteocytes on breast cancer cell proliferation in engineered two- and three-dimensional models. We observed that osteoblasts and osteocytes induce proliferative effects on cancer cells. Specifically, bone cells increase cancer proliferation in 2D culture and osteoblasts increase cancer growth more than osteocytes in 3D models. Moreover, osteocyte interaction with cancer cells in 3D models are stiffness dependent. We show that these effects depend on the CXCL5/CXCR2 signaling axis. Taken together, we demonstrate that osteoblasts drive cancer growth in a bone metastatic niche and that this effect can be rescued with CXCL5/CXCR2 inhibition.

## Introduction

Bone is the most common site for breast cancer metastasis, whereby bone metastases are present in ∼80% of patients with diagnosed metastasis and for over 90% of patients who succumb to the disease (1). Bone metastases are incurable and less responsive to standard treatments than primary tumors (2),. As such, patients with bone metastases live only 1.6 years after diagnosis on average (3). Beyond mortality, bone metastases are associated with morbidities such as irregular hematopoiesis, altered immune response, hypocalcemia or hypercalcemia, and disruption of bone turnover and maintenance leading to bone loss and pathological fractures (4). Current treatments for bone metastasis include radiation, surgery, and chemotherapy (5). Anti-resorptive drugs are often prescribed prophylactically at diagnosis of the primary tumor or following metastasis to reduce bone loss and its associated morbidities (5,6). Anti-resorptive drugs reduce tumor recurrence and improve survival in post-menopausal women, though do not show advantage in pre-menopausal women (7). A major challenge associated with breast cancer metastasis in the bone environment is that they can occur years or even decades after primary diagnosis and reported disease remission (8). Thus, tumor cells may enter a dormant state, which allows them to avoid diagnosis and treatment.

Breast cancer metastasis to bone is characterized by a vicious cycle in which cancer cells stimulate bone resorption, which releases biomolecules that support further cancer cell proliferation in a positive feedback loop (2,9–12). Within the bone marrow microenvironment, cancer cells secrete parathyroid hormone-related protein (PTHrP) and interleukins 6, 8, and 11 (IL-6, IL-8, and IL-11), which stimulate bone resorption by osteoclasts (9,13,14). The resulting release of sequestered transforming growth factor beta (TGF-β) from the bone matrix supports further tumor invasion leading to more bone resorption (1,2,15). Bone forming osteoblasts also support cancer progression by secreting chemokines including C-X-C motif ligands 5 and 12 (CXCL5, CXCL12) (4,16). In turn, tumor-secreted factors, such as DKK-1, compromise normal osteoblast activity, further contributing to the bone loss (17–19). Mechanical loading normally inhibits bone resorption and enhances formation through mechanosensitive osteocytes signaling to decrease osteoclast and increase osteoblast activity (1,20–24). However, loading also induces osteoblast expression of cytokines, such as CXCL1, CXCL2, and CXCL5, which contribute to metastatic progression by promoting invasion and metastasis and the transformation of adipocytes into cancer-associated adipocytes in the bone marrow (25).

Among the proteins and factors that support the breast cancer to bone vicious cycle, CXCL5, with its only known receptor CXCR2, promotes breast cancer colonization in bone by inducing escape from dormancy in disseminated tumor cells, which is likely responsible for late recurrence of metastatic disease (4,26–28) Indeed, inhibition of the CXCR2 receptor decreases proliferation, migration, and invasion of other cancers (27,29–31). The established role of the CXCL5/CXCR2 axis in breast cancer metastasis to bone suggests that it may be a druggable target for metastatic bone tumors. Indeed, chemokines are favorable targets to treat bone disease because they recruit immune cell populations (25). However, the bone cell population is diverse (32), and the source of CXCL5 and the interaction of the CXCL5/CXCR2 axis with other bone cell secreted proteins are not well understood.

Given the complex nature of the bone metastatic environment and breast cancer vicious cycle, *in vivo* or organ culture models, while robust, make it difficult to isolate roles of specific cell populations. Due to the mechanosensitive nature of osteoblasts and osteocytes and their unique morphology, 3D cultures can better replicate the bone microenvironment and support normal osteocyte morphology compared to 2D cultures (33–36). While various 3D models of breast cancer metastasis in bone aim to mimic *in vivo* bone matrix structures, recapitulating the interactions between bone and tumor remains a challenge (36–43). A bone mimetic model created by co-culturing osteocytes, osteoblasts, and pre-osteoclasts in gelatin hydrogels containing nano-hydroxyapatite (nano-HA) particles supports osteoblast mineralization, osteoclast differentiation, and formation of osteocyte cell processes (36,37,44). Gelatin encapsulation also allows cells to be spatially patterned to study specific cell interactions, and the mechanical properties can be modulated by chemical crosslinking. Together, these features provide a platform to characterize interactions of bone and cancer cells and the mechanical microenvironment on tumor growth in an intermediate model of bone metastasis (20,36).

The objective of this study was to determine whether CXCR2 inhibition between cancer and bone cells modulates cancer cell proliferation and tumor spheroid formation. We employed both a 2D multilayered model of the bone niche and a 3D co-culture model of osteoblasts, osteocytes, and breast cancer cells in gelatin hydrogels to study how secreted factors from osteocytes and osteoblasts modulate 4T1 triple negative breast cancer cell proliferation, tumor spheroid formation, and gene expression (44). Specifically, we 1) compared 4T1 cell proliferation, tumor spheroid size, and gene expression in the presence of osteoblasts, osteocytes, and their combination; 2) determined the effects of matrix stiffness on the interactions between bone and cancer cells; and 3) quantified the effects of CXCL5/CXCR2 axis inhibition on cancer cells proliferation, spheroid size, and/or gene expression in the presence of bone cells.

## Materials and Methods

### Cells and Reagents

Murine OCY454 osteocytic cells (Centre for Skeletal Research Bone Cell Core, Massachusetts General Hospital) were cultured at the permissive temperature of 33°C for two weeks on type 1 collagen (0.15 mg/ml in 0.02 M acetic acid) coated tissue culture flasks. Cells were then transferred to non-coated flasks and, after 3 days, differentiated by culturing at the semi-permissible temperature of 37 °C for two weeks. Differentiated OCY454 cells were expanded in α-MEM (Millipore Sigma) supplemented with 10% Fetal Bovine Serum (FBS, Corning), 2% antibiotic antimycotic solution (ABAM, Corning), and 2 mM L-glutamine.

Murine MC3T3-E1 osteoblastic cells (ATCC, Manassas, VA, USA) were cultured under standard conditions (5% CO_2_, 37°C) in supplemented α-MEM. Murine 4T1 metastatic mammary carcinoma cells were cultured under standard conditions in RPMI 1640 culture media (ThermoFisher) containing 10% FBS and 2% ABAM. On the day of experiment, 4T1 cells were stained with PKH26 red fluorescent cell linker for general cell membrane labeling (Sigma-Aldrich MIDI26), following the manufacturer’s protocol. Briefly, suspended cells were washed with serum-free media. A dye solution was incubated for 5 minutes. Following incubation, serum was added to stop the staining, and the cells were centrifuged and washed three times in phosphate-buffered saline (PBS).

### Transwell Experiments

Transwell inserts were used to recreate a simple bone-like niche (Fig. 1) (45). Osteoblasts and osteocytes were cultured on opposite sides of transwell cell culture plate inserts containing permeable membranes (Corning, 3 µm pores). The pore size was chosen to allow the OCY454 and MC3T3-E1 cells to form functional gap junctions across the membrane (46). Briefly, inserts were first inverted and 2,600 MC3T3-E1 cells/cm^2^ were seeded on the basal surface of the insert in 500 μL of medium and allowed to attach for 6 hours. The inserts were then placed right side up into 12 well plates and 2,600 OCY454 cells/cm^2^ were seeded on the apical side of the insert and 2,600 4T1 cells/cm^2^ were seeded in the bottom of the well. We further investigated the role of OCY454 or MC3T3 cells separately by seeding them individually on the apical sides of the membranes with 4T1 cells seeded on the bottom of the culture plates all in the same media used for the 3D culture (Table 1).

**Figure 1:**
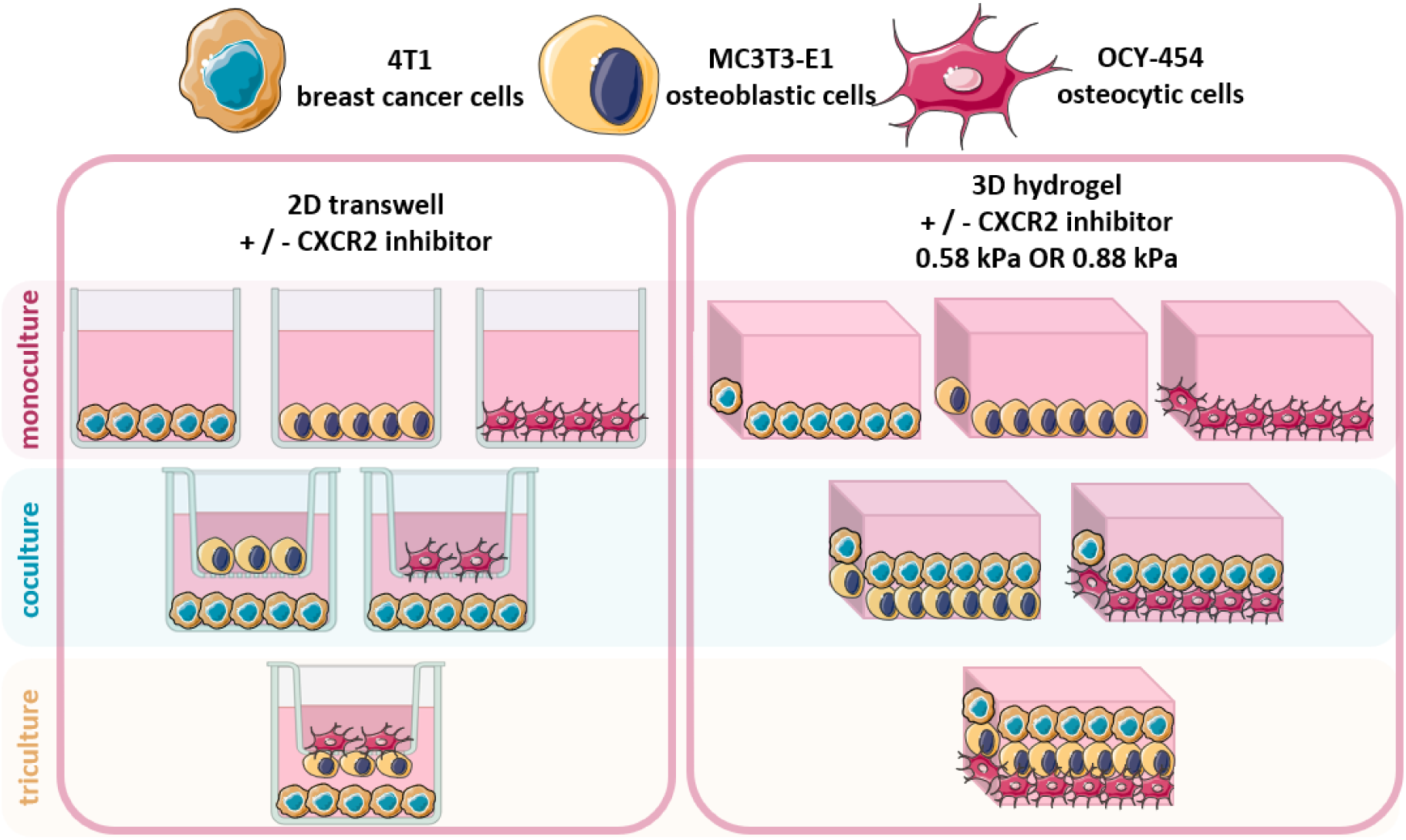
Experimental design schematic. 4T1 metastatic breast cancer cells, MC3T3-E1 osteoblastic cells, and OCY454 osteocytic cells are seeded either on transwell cell culture plates or in hydrogels of different stiffness (0.58, 0.88 kPa) in monoculture, co-culture, and tri-culture configurations.

**Table 1.** Hydrogel culture media formulations.

| Culture Condition | Media Formulation |
| --- | --- |
| Monoculture | 4T1 growth media |
| Osteoblast co-culture | 50% 4T1 growth media, 50% MC3T3/OCY454 growth media |
| Osteocyte co-culture | 50% 4T1 growth media, 50% MC3T3/OCY454 growth media |
| Tri-culture | 33% 4T1 growth media, 66% MC3T3/OCY454 growth media |

Proliferation of 4T1 cells was quantified on days 1, 3, 5, and 7 using the CellTiter-Blue Cell Viability Assay (Promega) following manufacturers protocols. Briefly, membrane inserts were removed and temporarily stored in a 12 well plate with appropriate media. CellTiter-Blue was added to media at a 1:5 ratio and allowed to incubate for 1 hour. Fluorescence was measured using a plate reader and normalized to known 4T1 concentrations.

### Hydrogel Fabrication

To investigate the effects of osteoblast and osteocyte paracrine interactions on cancer cells, cancer and bone cells were cultured in gelatin-transglutaminase hydrogels, doped with nano-HA, in different combinations: (1) cancer monoculture (4T1), (2) cancer-osteoblast co-culture (4T1+MC3), (3) cancer-osteocyte co-culture (4T1+OCY), and (4) tri-culture (4T1+MC3+OCY) (Fig. 1) using methods adapted from Kumar et al., 2024 (44). Briefly, gelatin (Type A, 175 bloom Sigma) was mixed in supplemented RPMI media at a concentration of 90 mg/mL, heated for 30 minutes at 60 °C, sterile filtered using a 0.2 μm filter, and stored for 24 hours at 4 °C. Transglutaminase (Activa WM; Ajinomoto foods Europe S.A.S., France) was mixed with supplemented RPMI media at concentrations of 0.3% or 0.8% per gram of gelatin (27 or 72 mg/mL) to yield low- and high-stiffness gels, respectively (37,43). Transglutaminase solutions were sterile filtered using a 0.2 μm filter, and stored for 24 hours at 4 °C. Nano-hydroxyapatite (nano-HA) particles were created on the day of experiment by mixing a 0.02 M calcium chloride dihydrate solution (2.92 mg/mL calcium chloride dihydrate in ultrapure water) with a phosphate solution (50mM HEPES buffer, 140 mM sodium chloride, 5 mM sodium hydroxide, 0.012 M sodium phosphate tribasic dodecahydrate, 0.017% v/v Darvan 821A in DI water) at a 1:2 ratio with DI water and sterile filtered using a 0.2 μm filter.

Before fabrication, cells were counted and suspended in media to obtain cell density of 2×10^6^ cells/ mL. The cell suspension was mixed 1:1 with the nano-HA solution prior to fabrication. To fabricate hydrogels, the gelatin solution was heated to 37°C and mixed with the transglutaminase solution and cell suspension/nano-HA solution at a 1:1:1 ratio and pipetted into custom-made polydimethylsiloxane (PDMS) molds (1 mm D × 4 mm W × 13 mm L) and allowed to crosslink at 4 °C for 8 min. Hydrogel layers containing different cell types were formed on top of the construct after each layer had crosslinked. Mono- and co-culture constructs included cell-free gelatin layers so that all constructs had the same depth, see Figure 1. The constructs were cultured with different media formulations (Table 1) for 7 days. In the treated groups, media was supplemented daily for seven days with SB225002 (MedChemExpress, 100 nM final concentration per well), a nonpeptide inhibitor and antagonist of CXCR2 (47).

Following 7 days of culture, half of the constructs from each group were fixed in 4% paraformaldehyde (PFA) for 24 hours at 4 °C. Fixed constructs were washed three times in PBS and stored in PBS at 4°C for immunofluorescence staining. The remaining hydrogels were removed from culture on day 7 and immediately frozen at −80°C for PCR.

### Immunohistochemistry

The fixed gels were immunostained for Ki67 (AB9260, Sigma), actin (phalloidin-iFluor488, AAT Bioquest), and nuclei (DAPI dilactate, Millipore Sigma). The cells were permeabilized within the constructs by incubating in 0.5% Triton-X in PBS for 10 min at 4°C. The gels were washed in BSA (Sigma A7906) three times and then incubated in BSA for 1 hour at room temperature.

For Ki-67 staining, constructs were incubated overnight at 4°C in 150 mL of the antibody solution, diluted at 1:200 for rabbit anti-Ki67 primary antibody (AB9260, Sigma). After washing three times with BSA for 10 minutes at room temperature, the Ki67 stained gels were incubated in the secondary antibody solution, donkey anti-rabbit FITC (711-095-152, Jackson ImmunoResearch), for 90 minutes at room temperate on a shaker.

Finally, after three washing cycles, constructs were incubated in phalloidin-FITC (1:400) to stain the actin cytoskeleton (for gels not stained with Ki67) and DAPI dilactate (1:2000) (for all gels) to stain the nucleus for 30 minutes at room temperature and rinsed again with 1% BSA solution. Constructs were stored in PBS at 4°C until fluorescence was detected using confocal laser scanning microscopy (Olympus Fluoview 3000).

### Quantitative Real-Time PCR

Samples frozen on day 7 were thawed, and total RNA isolation was performed. Constructs were placed in 1 mL of Trizol™ Reagent (Invitrogen) and homogenized using a 16-gauge needle and syringe. 200 μL of chloroform was added and the solution was centrifuged at 12000 × *g* separating out a clear aqueous supernatant. The clear aqueous phase was removed using a pipette and purified using RNEasy kit (Qiagen, Germantown, PA) according to the manufacturer’s instructions. RNA yield was measured using the DS-11 FX (DeNovix). 500 ng of RNA was converted to cDNA using the Qiagen Quantitect Reverse Transcription Kit, following manufacturers protocol, and thermal cycler (5PRIMEG/O2, Prime). Relative gene expression was investigated by quantitative real-time polymerase chain reaction (qRT-PCR). Primers for sclerostin (SOST), receptor activator for nuclear factor κ B Ligand (RANKL), osteoprotegerin (OPG), osteopontin (OPN), parathyroid hormone related protein (PTHrP), and CXC chemokine motif ligand 5 (CXCL5) were used (Table 2). RPLPO was used as a reference gene. qRT-PCR was completed with 25 ng of cDNA per well using Qiagen Quantitect SYBR Green PCR Kit and a Quant Studio 5 PCR machine (Applied Biosciences). Differential gene expression was analysed using the δ-δ CT method to calculate the fold-change in expression relative to the corresponding low stiffness or untreated condition.

**Table 2.**
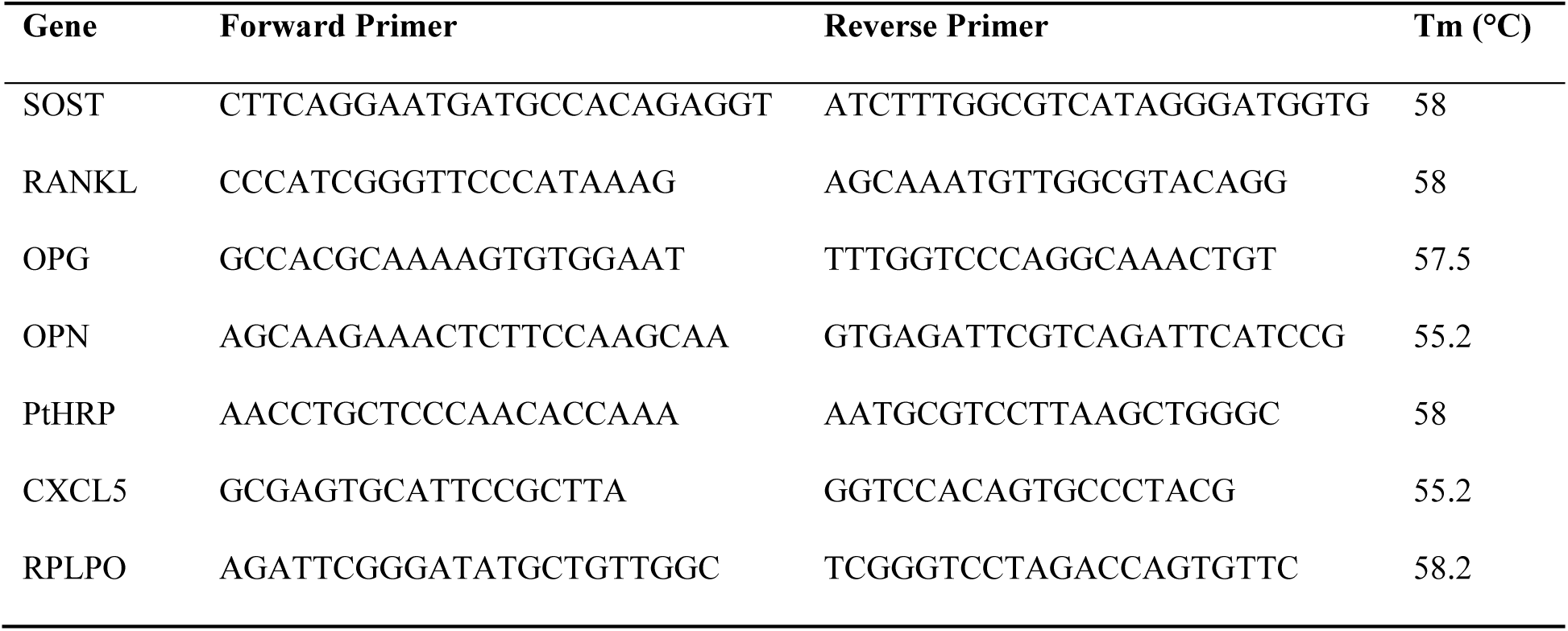
Primers (5’-3’) and annealing temperatures (Tm) employed for qRT-PCR.

### Statistical Analysis

Statistical analysis was performed on GraphPad Prism (version 8) software. Data sets were tested for normality and lognormality, and Kruskal-Wallace one factor ANOVA was used to compare groups for spheroid diameter and Ki67 intensity measurements. One sample t-tests with a null hypothesis that the mean = 1 were used to compare qRT-PCR data. Results are displayed as mean ± standard deviation. Significance was accepted at p ≤ 0.05.

## Results

We investigated the impact of bone osteoblasts and osteocytes on cancer cell proliferation and tumor spheroid formation using both transwell and 3D co-cultures. The murine 4T1 mammary cancer cells were co-cultured with combinations of murine MC3T3-E1 osteoblastic cells and murine OCY454 osteocytic cells. Samples were analyzed for spheroid diameter, Ki67 expression as a marker of cell proliferation, and gene expression of PTHrP and CXCL5. We also tested the impact of matrix stiffness on these factors by comparing these parameters on cells grown in 3D on hydrogels with soft or stiff matrix.

### Transwell co-culture of cancer cells with bone cells increases cancer cell proliferation

In our 2D transwell experiments, the cancer cell count increased in osteoblast co-culture, osteocyte co-culture, and tri-culture when compared to 4T1 monoculture groups at 5 days (Fig. 2A; p < 0.01). Tri-culture cancer cell counts were lower than osteocyte co-culture (p < 0.05).

**Figure 2:**
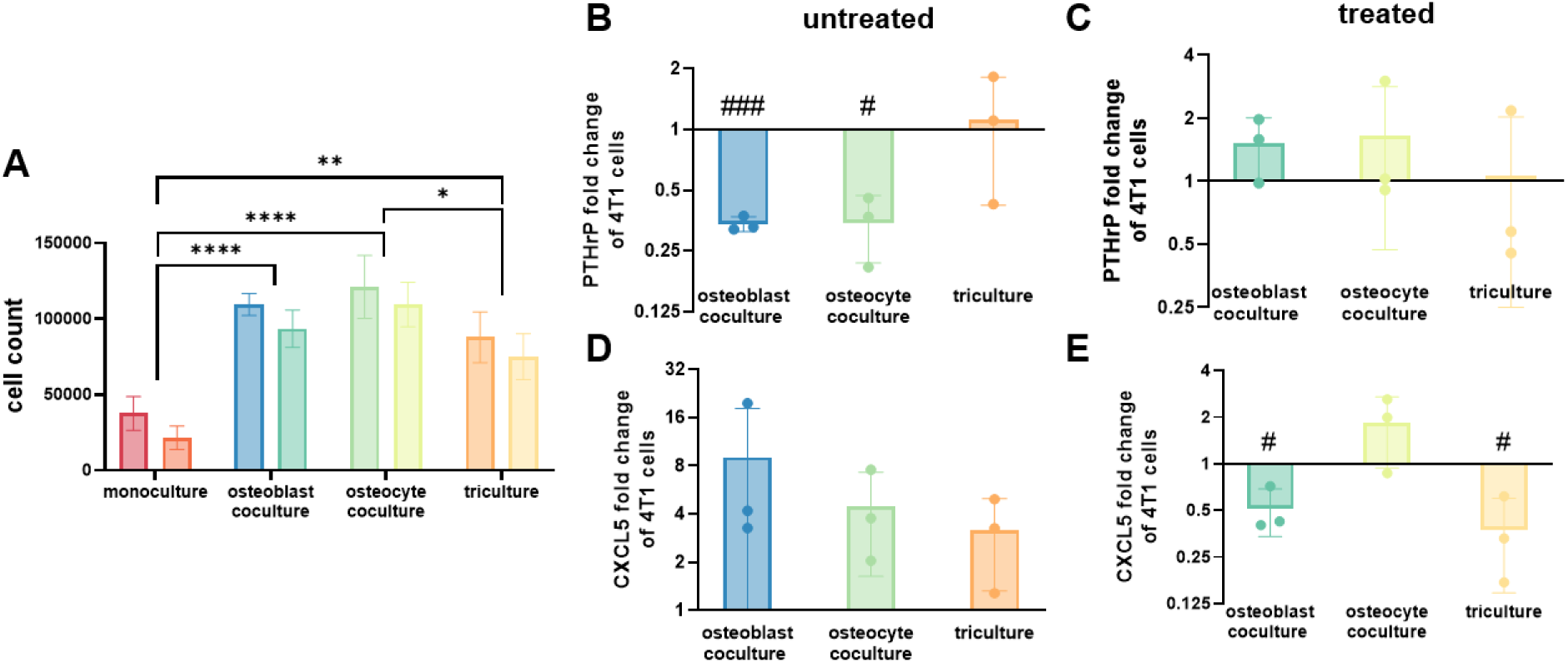
Transwell co-culture with bone cells increases cancer cell proliferation. **A)** CellTiter-Blue assay on day 5 of culture revealed bone cell co-culture increased 4T1 cancer proliferation in transwell experiments. **(B-C)** PTHrP expression by qRT-PCR. Fold change of 4T1 breast cancer gene expression for PTHrP downregulated in osteoblast and osteocyte co-culture in **(B)** untreated conditions normalized to monoculture and remained unchanged in **(C)** treated normalized to untreated conditions. **(D-E)** CXCL5 expression by qRT-PCR. Fold change of 4T1 breast cancer gene expression for CXCL5 remained unchanged in **(D)** untreated conditions normalized to monoculture and downregulated in osteoblast co-culture and tri-culture groups in **(E)** treated normalized to untreated conditions. (n=3, # p < 0.05 vs 1.0, ### p < 0.001) (* p < 0.05; ** p < 0.01; *** p < 0.001; **** p 0.0001).

To test for the requirement for CXCR2 in these cultures, the cell cultures were treated with or without SB225002, which is a CXCR2 antagonist (antiCXCR2). While antiCXCR2 treatment did not significantly affect cancer cell count in 2D, it consistently decreased cell count in all groups (Fig. 2A).

PTHrP is a protein secreted by cancer cells that stimulates bone resorption by osteoclasts. Osteoblast and osteocyte co-culture downregulated PTHrP gene expression in 4T1 cells compared to 4T1 monoculture (Fig. 2B; p < 0.05), whereas tri-culture had no effect. CXCR2 inhibition did not significantly affect PTHrP gene expression in 4T1 cells compared to untreated groups (Fig. 2C).

Bone cell co-culture had no effect on CXCL5 gene expression by qRT-PCR in untreated groups normalized to monoculture (Fig. 2D). However, the presence of osteoblasts in osteoblast co-culture and tri-culture reduced CXCL5 expression in antiCXCR2 groups normalized to untreated conditions (Fig. 2E; p < 0.05).

### Osteoblasts increase cancer cell proliferation in 3D co-culture, whereas osteocytes effects are dependent on substrate stiffness

Cancer cells formed spheroids in all hydrogel constructs by 3 days (Supplemental Fig. 1A), consistent with previous studies (43). Overall, spheroid sizes were larger in soft gels compared to stiff. In soft cultures (0.5 kPa), osteoblast co-cultures formed larger 4T1 spheroids when compared to monoculture, osteocyte co-culture, and tri-culture groups (Fig. 3B; p < 0.01). In stiff hydrogels, the osteoblast and osteocyte co-culture increased the 4T1 spheroid diameter compared to monoculture and tri-culture groups (Fig. 3C; p < 0.05).

**Figure 3:**
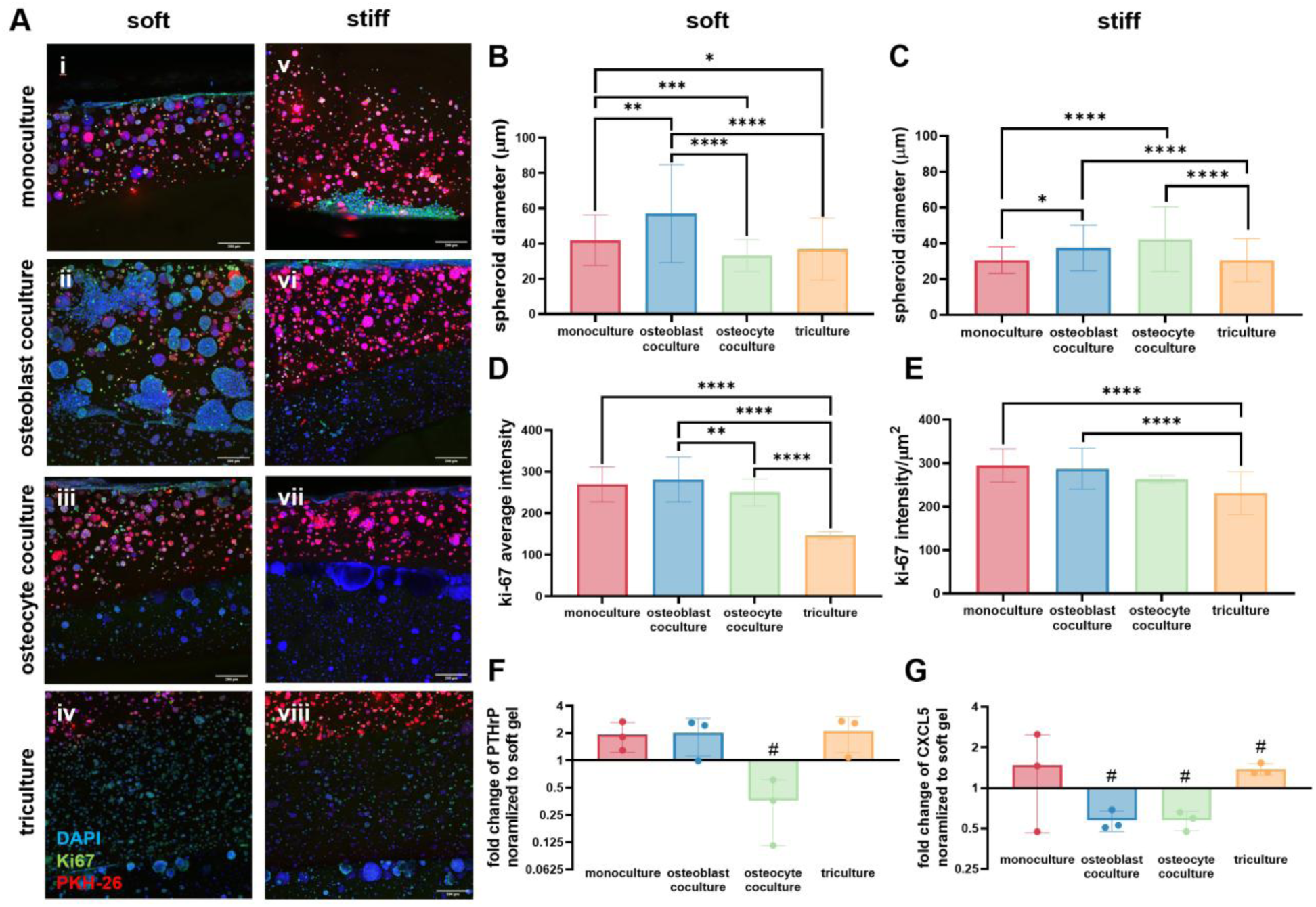
Osteoblasts increase cancer cell proliferation in 3D culture, whereas osteocyte effects are dependent on substrate stiffness. **(A)** Immunofluorescence. Representative immunofluorescence-stained hydrogels for DAPI (blue), Ki67 (green), and PKH26 (red) at day 7 (scale bar: 200 μm). **(B-C)** Spheroid diameter in **(B)** soft and **(C)** stiff gels. In soft gels, osteoblast co-culture increased day 7 4T1 spheroid diameter compared to monoculture, osteocyte co-culture, and tri-culture. In stiff gels, osteoblast and osteocyte co-culture increased 4T1 spheroid diameter compared to monoculture and tri-culture. **(D-E)** Ki67 expression in **(D)** soft and **(E)** stiff gels. In soft gels, monoculture and osteoblast co-culture increased day 7 Ki67 intensity compared to osteocyte co-culture and tri-culture. **In stiff gels,** tri-culture decreased Ki67 intensity compared to monoculture and osteoblast co-culture. (* p < 0.05; ** p < 0.01; *** p < 0.001; **** p 0.0001). **F-G)** PTHrP expression in **(F)** soft and **(G)** stiff gels by qRT-PCR. Fold change of PTHrP gene expression decreased in stiff osteocyte co-culture and was unchanged in other groups normalized to respective soft conditions. Fold change of CXCL5 gene expression decreased in stiff osteoblast and osteocyte co-cultures and increased in tri-culture normalized to respective soft conditions (n=3; # p < 0.05 vs 1.0).

Ki67 is a marker of cell proliferation. Ki67 intensity on day 3 was higher in osteoblast co-culture compared to other conditions (Supplemental Fig. 1B; p < 0.0001). In contrast, Ki67 intensity decreased in soft and stiff tri-cultures compared to other conditions on day 7 (Fig. 3D,E; p < 0.01). Because Ki67 intensity shows cells in active proliferation (48), the proliferation at day 3 may explain the increased spheroid diameter of the osteoblast co-culture groups by day 7.

Stiff osteocyte co-culture downregulated PTHrP gene expression when compared to soft osteocyte co-culture (Fig. 3F; p < 0.05). Stiffness did not affect PTHrP expression in any other culture conditions.

CXCL5 expression by qRT-PCR decreased for both stiff osteoblast and osteocyte co-cultures but increased by stiff tri-culture and was unchanged in monoculture (Fig. 3G; p < 0.05).

### CXCR2 inhibition abrogates osteoblasts pro-proliferative effect on cancer cells

Taking a closer look at our soft gels, in which larger cancer spheroids were formed, the addition of the CXCR2 antagonist in osteoblast co-culture decreased cancer spheroid size (p < 0.0001) to similar sizes as monoculture (Fig. 4B; p > 0.05). Consistently, Ki67 intensity decreased in osteoblast co-cultures with antiCXCR2 treatment, while intensity was unchanged in all other groups (Fig. 4B; p < 0.05). Treatment with antiCXCR2 also decreased spheroid size in tri-culture (p < 0.01), while monoculture spheroid diameter was unaffected. Osteocyte co-culture, which exhibited smaller cancer spheroids than monoculture in soft gels, had larger spheroids with antiCXCR2 treatment (p < 0.0001), similar to monoculture (p > 0.05).

**Figure 4:**
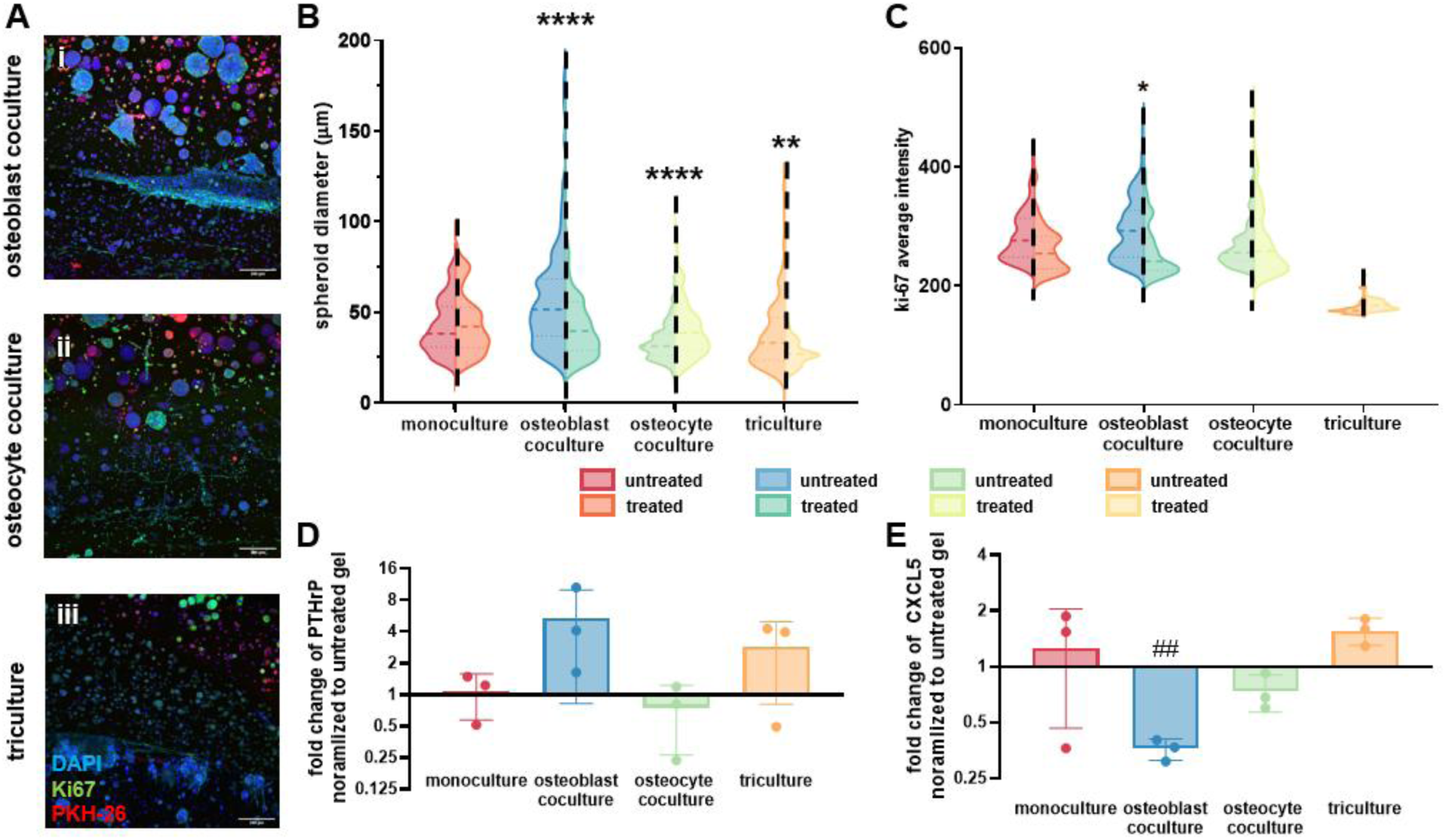
CXCR2 inhibition abrogates osteoblasts pro-proliferative effect on cancer cells. **A)** Immunofluorescence. Representative immunofluorescence-stained for DAPI (blue), Ki67 (green), and PKH26 (red) of soft CXCR2 antagonist treated **(i)** osteoblast co-cultures, **(ii)** osteocyte co-cultures, and **(iii)** tri-cultures at day 7 (scale bar: 200 um). **B)** Spheroid diameter. AntiCXCR2 treatment reduced 4T1 spheroid size in osteoblast co-culture and tri-culture groups, increased spheroid size in osteocyte co-culture groups, and had no effect in monoculture. **C)** Ki67 expression. AntiCXCR2 treatment reduced Ki67 intensity at day 7 in osteoblast co-culture groups and had no effect in monoculture, osteocyte co-culture, and tri-culture. (* p < 0.05; ** p < 0.01; *** p < 0.001; **** p < 0.0001). **(D-E)** Gene expression by qRT-PCR. Fold change of **(D)** PTHrP and **(E)** CXCL5 gene expression normalized to respective untreated gels (n=3; ## p < 0.01).

When comparing treated to untreated gene expression by qRT-PCR, PTHrP was unchanged by antiCXCR2 treatment (Fig. 4D). CXCR2 antagonist treatment in osteoblast co-culture downregulated CXCL5 gene expression but exhibited no effect in other culture conditions (Fig. 4E; p < 0.01). CXCR2 inhibition increased PTHrP and CXCL5 gene expression in several culture conditions in stiff gels (Supplemental Figure 2).

## Discussion

Metastatic breast cancer in bone is highly aggressive and is associated with comorbidities and increased mortality rates, necessitating an improved understanding of this disease and novel treatments to combat its progression (3,8). We used novel *in vitro* models in 2D and 3D co-cultures to study the paracrine interactions between osteoblasts, osteocytes, and breast cancer cells. Our goals were to determine how osteoblasts and osteocytes affect cancer cell proliferation and spheroid growth and investigate whether this was mediated by CXCR2 ligands. We found that bone cells increase cancer cell proliferation and alter metastatic markers in 2D and 3D cultures. Specifically, we revealed in this model that osteoblasts increase breast cancer cell growth more than osteocytes and in a CXCR2 dependent manner. We also found that osteocytes alter cancer growth in 3D cultures differently in different stiffnesses.

We found that breast cancer spheroid growth was enhanced by osteoblasts in a CXCL5/CXCR2 dependent manner, as CXCR2 inhibition rescued the increase in both cancer spheroid growth and Ki67 intensity in our soft hydrogel constructs. In osteoblast co-culture, CXCL5 gene expression decreased after treatment with antiCXCR2. Though osteoblast-induced cancer proliferation was not rescued by CXCR2 inhibition in 2D transwell cultures, the gene expression of these cultures did reveal that with antiCXCR2, the presence of osteoblasts in these cultures downregulated CXCL5 expression, both in co-culture and tri-culture. This is consistent with several studies that have shown that CXCL5 signaling is implicated in bone metastasis (4,27,49), though few studies have shown that the role of CXCL5 in bone metastases is osteoblast dependent (16). Very few studies have investigated osteocyte CXCL5/CXCR2 signaling (50), and no studies have shown their specific involvement in this pathway during breast cancer. Our results indicate that osteocytes are not highly implicated in the CXCL5/CXCR2 signaling pathway, though more physiologically relevant models of osteocytes and metastasis are warranted to confirm this. While osteoblasts are heavily implicated in the breast cancer vicious cycle (2,16,17,51–57), they have not been specifically shown to drive tumor progression. Our results indicate that osteoblasts, more so than osteocytes, have a pro-tumorigenic effect on cancer cells, especially in 3D mimetic cultures.

Our study provides a powerful platform to study the interactions between bone and cancer cells in 2D and 3D cultures. Importantly, the 3D cultures capture the complex microenvironment of breast cancer metastases in bone, with our matrix stiffnesses being similar to bone marrow *in vivo* (58). We found that the mechanical environment alters the interactions between cancer and bone cells, both changing how the cancer spheroids grow and how osteoblast and osteocyte paracrine factors affect cancer growth. Our findings suggest that CXCL5/CXCR2 signaling plays a significant role in cancer cell interactions within the bone microenvironment, uncovering that osteoblasts have a greater influence on the cancer cells than the osteocytes.

Several limitations need to be considered in this study. First, the stiffness of our hydrogels are about six orders of magnitude lower than the stiffness of bone. As such, osteocytes were not in normal physiological conditions, which may have affected their physiology. Indeed, we demonstrated that osteocyte behavior depended on the gelatin stiffness in this model. Conversely, the gelatin stiffness is a reasonable approximation for bone marrow (58), which is appropriate for the cancer cells. Second, our experimental timelines were too short to allow the hydrogels to fully mineralize (44). This may have affected the bone cell proteome and interactions with cancer cells, though shorter experimental timelines provide insight into early interactions between cancer and bone cells. Third, our gene expression measures were conducted using pooled cell populations in the hydrogels rather than separated by cell type. Thus, the reported differences in gene expression cannot be attributed to specific cell populations. Gene expression we did quantify in individual cell populations in our transwell plates, which provides insight into their potential.

Here we reveal that both 2D and 3D co-culture of breast cancer cells with osteoblasts, osteocytes, or both increases cancer cell proliferation. In 2D culture, the effects did not differ between bone cell types, although tri-culture conditions induced less cancer cell proliferation than co-culture. This may be due to the increase in competition for nutrients by the cells in tri-culture. In contrast, 3D culture indicated that osteoblasts consistently increase cancer cell proliferation, while osteocyte and tri-culture effects depended on the local mechanical environment. This may be due to the effects that 3D culture have on osteocyte behavior. Osteocytes are well established as mechanically sensitive cells that respond to and secrete protein and small molecules that are regulated by their mechanical environment (34,35,59). Capturing osteocytes *in vivo* is challenging due to these cells normally being embedded in the stiff bone matrix. While 3D culture can support the development of the normal dendritic morphology of osteocytes (34,36,60), hydrogels are several orders of magnitudes softer than bone. Thus, neither the 2D nor 3D cultures can fully capture osteocyte interaction with cancer cells. As osteocyte effects on cancer proliferation are inconsistent in 3D cultures, how the mechanical environment of osteocytes affects their behavior and interaction with cancer cells must be further studied. Taken together, our results indicate that metastatic breast cancer’s affinity for bone and growth in the bone microenvironment may be primarily driven by osteoblasts through CXCR2/CXCL5 signaling while osteocytes have a context dependent effect.

Co-culture with bone cells in 2D transwell plates downregulate PTHrP gene expression in 4T1 cells, especially in osteoblast co-culture. The reported role of PTHrP in breast cancer progression is variable (61), with several studies indicating PTHrP expression in cancer cells improves prognosis and cancer progression and reduces bone metastases (62–64), while others indicate that PTHrP increases bone metastases and cancer cell invasion (65–67). PTHrP is also expressed by osteoblasts (68,69), and in the bone metastatic environment plays a role in paracrine, autocrine, intracrine and endocrine signaling (61). The results in this study indicate that PTHrP is implicated in promoting cancer growth, and that the expression of PTHrP is not dependent on CXCL5/CXCR2 signaling. However, the complex effects of this protein and its related pathways must be further studied to comprehensively understand its role in bone metastases.

Our data complements previous studies where osteoblast-derived factors increased breast cancer cell migration (16,53). We expanded on these studies by comparing osteoblast and osteocyte effects on cancer cells. Osteocyte secreted factors have differing effects on cancer progression based on cancer type (70), with osteocytes having inhibitory effects on breast cancer growth in some cases (71,72). However, we found this inhibitory effect was mechanically regulated with osteocyte co-culture eliciting larger cancer spheroids than monoculture in stiffer matrices.

Taken together, our novel model of breast cancer metastasis to bone uncovered that osteoblasts have a pro-proliferative effect on cancer cells that is CXCR2 dependent, while osteocytes have a mechanical environment dependent effect that is independent of CXCR2 activation.

## Supporting information

Raw Data

## Acknowledgements

SN was supported by the Harper Cancer Research Institute Interdisciplinary Interface Training Program (IITP) Grant through the Walther Cancer Foundation. This research was supported through GLN, LEL, and LMM by the Notre Dame Naughton Accelerator Grant, the U.S. National Institute of Health (NIH) RO1 CA252878, and the Department of Defence (DOD) Breast Cancer Research Program (BCRP) Breakthrough Award, Level 2 (W81XWH2110432), the Irish Research Council Laureate Award Programme 2017/18 (MEMETic, IRCLA/2017/217), and the European Research Council (ERC) Consolidator grant (MEMETic 863795). The authors acknowledge the assistance and facilities of the Centre for Microscopy and Imaging at the University of Galway.

## Author Contributions

SN performed the main experiments, data curation and analysis, and drafted the manuscript. SMN assisted with project conception, experiments, and data curation. IW and NV assisted with experiments. VK assisted with methodology. LEL supervised, assisted with data presentation, and secured funding. GLN and LMM conceived and supervised the project, edited the manuscript, and secured funding. All authors reviewed the final manuscript.

## Supplemental Figures

**Supplemental Figure 1:**
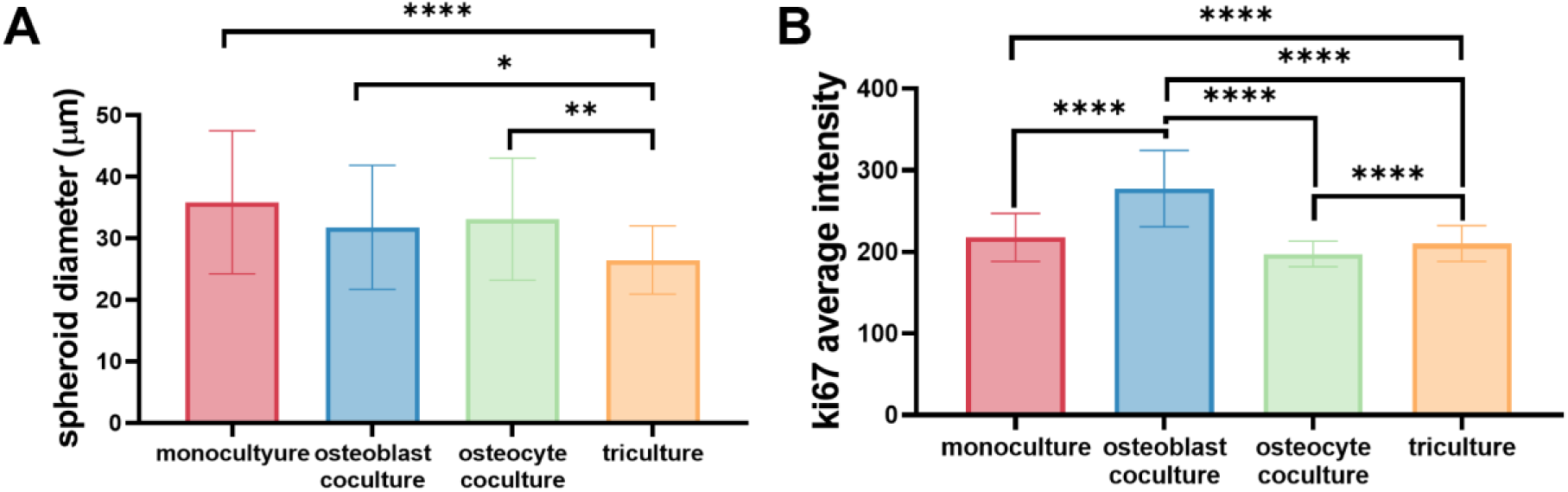
Day 3 4T1 spheroid growth. **A)** Spheroid diameter. Tri-culture reduced 4T1 spheroid diameter compared to monoculture, osteoblast co-culture, and osteocyte co-culture groups. **B)** Ki67 expression. Osteoblast co-culture increased Ki67 intensity compared to monoculture, osteocyte co-culture, and tri-culture groups (* p < 0.05; ** p < 0.01; **** p < 0.0001).

**Supplemental Figure 2:**
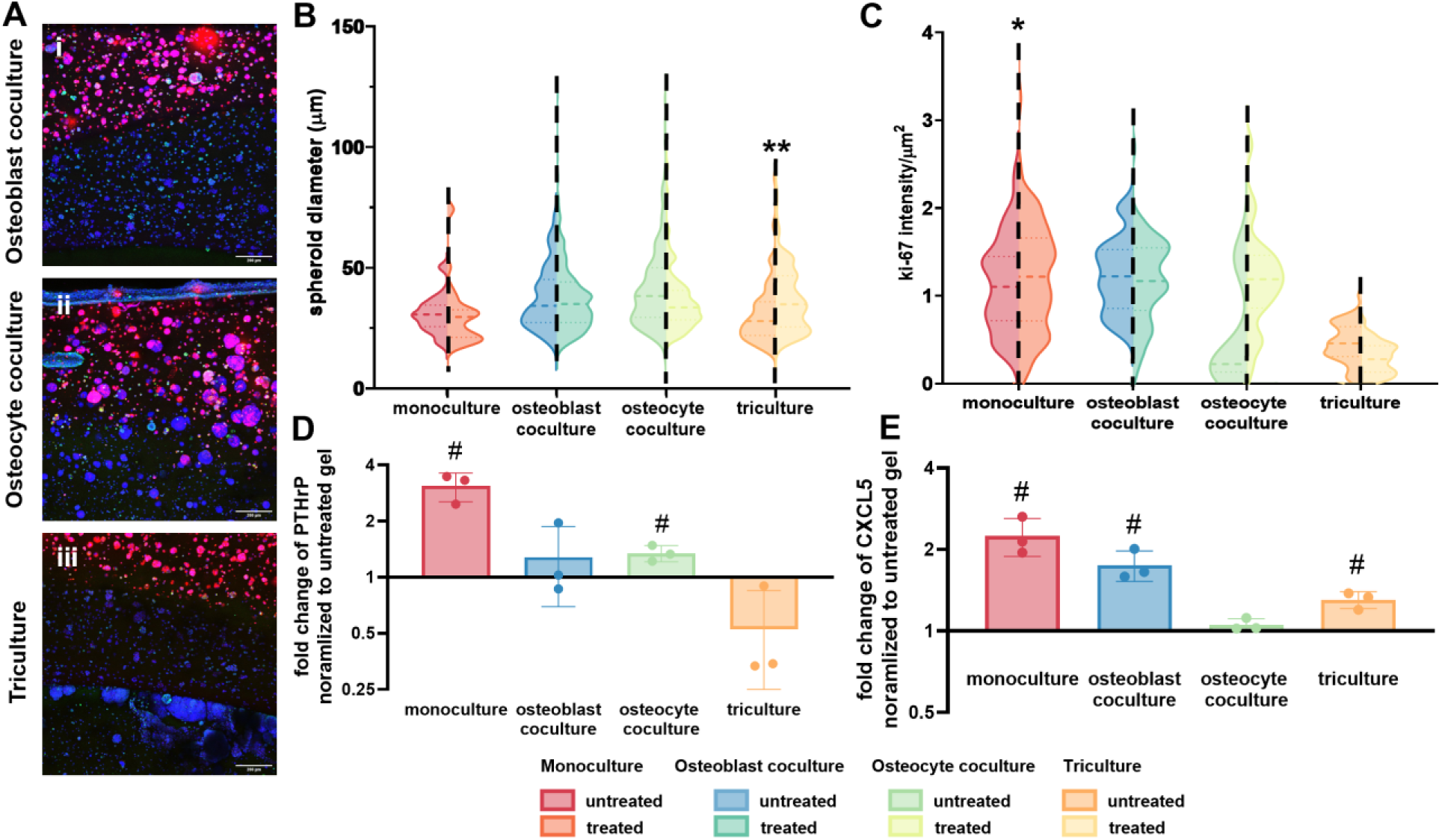
Effects of CXCR2 inhibition on 4T1 spheroid growth in stiff gels. **A)**Immunofluorescence. Representative immunofluorescence stained for DAPI (blue), Ki67 (green), and PKH26 (red) of stiff CXCR2 antagonist treated **(i)** osteoblast co-cultures, **(ii)** osteocyte co-cultures, and **(iii)** tri-cultures at day 7 (scale bar: 200 um). **B)** Spheroid diameter. AntiCXCR2 treatment increased 4T1 spheroid size in tri-culture groups and had no effect in other culture conditions. **C)** Ki67 expression. AntiCXCR2 treatment increased Ki67 intensity at day 7 in monoculture and remained unchanged in other groups. (* p < 0.05; ** p < 0.01). **(D-E)** Gene expression by qRT-PCR. Fold change of **(D)** PTHrP and **(E)** CXCL5 normalized to respective untreated gels (n=3; # p < 0.05).

