## Supplemental Figures for "Osteoblasts Exert a Pro-Tumorigenic Effect on Breast Cancer Spheroids Through CXCL5/CXCR2 Signaling In 2D And 3D Bone Mimetic Cultures"

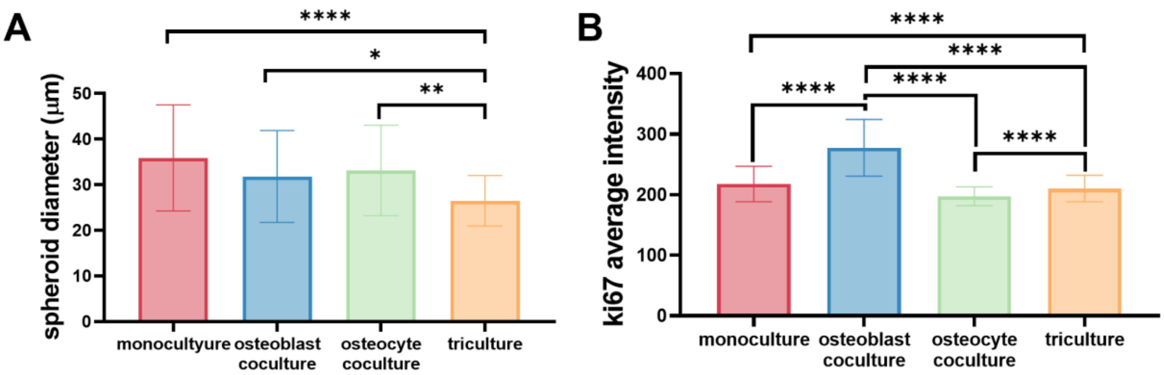


**Supplemental Figure 1: Day 3 4T1 spheroid growth. A)** Tri-culture reduced 4T1 spheroid diameter compared to monoculture, osteoblast co-culture, and osteocyte co-culture groups. **B)** Osteoblast co-culture increased ki67 intensity compared to monoculture, osteocyte co-culture, and tri-culture groups (* $p < 0.05$; ** $p < 0.01$; **** $p < 0.0001$).

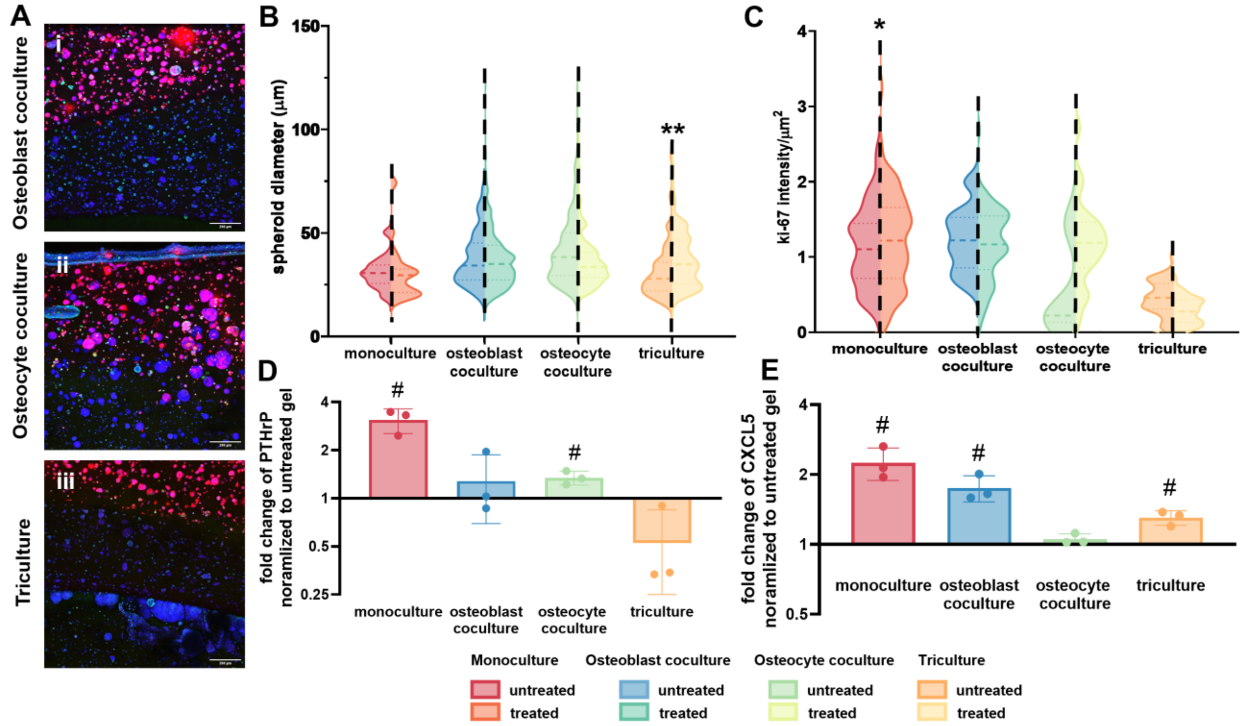


**Supplemental Figure 2: Effects of CXCR2 inhibition on 4T1 spheroid growth in stiff gels.** **A)** Representative immunofluorescence-stained for DAPI (blue), Ki67 (green), and PKH26 (red) of stiff CXCR2 antagonist treated **(i)** osteoblast co-cultures, **(ii)** osteocyte co-cultures, and **(iii)** tri-cultures at day 7 (scale bar: 200 um). **B)** antiCXCR2 treatment increased 4T1 spheroid size in tri-culture groups and had no effect in other culture conditions. **C)** antiCXCR2 treatment increased Ki67 intensity at day 7 in monoculture and remained unchanged in other groups. (* $p < 0.05$; ** $p < 0.01$). Fold change of **(D)** PTHrP and **(E)** CXCL5 normalized to respective untreated gels (n=3; # $p < 0.05$).
